# TKTDsimulation.jl and tktdjl2r: innovative packages for High Performance Computing of survival predictions in support of environmental risk assessment under time-variable scenarios

**DOI:** 10.1101/2021.02.18.431769

**Authors:** Virgile Baudrot, Sandrine Charles

## Abstract

Predictive environmental risk scenarios are today of major interest for environmental risk assessment as they provide plausible and consistent descriptions of possible effects of chemical *in natura*. In particular, they can be used for predictions of the future as consistent descriptions of pathways towards desired targets to protect. One single scenario would therefore be meaningless, as it could not capture all the variability and uncertainty involved in natural phenomenon combined with socio-economical events. A set of environmental risk scenarios is then a key asset to address sustainable and collaborative decision making associated with appropriate actions.

Toxicokinetics-Toxicodynamics (TKTD) models are increasingly used for the assessment and the prediction of environmental risk assessment due to chemical products. This mechanistic modelling approach offers many advantages as the possibility to perform simulations under non-observed realistic situations with time-variable exposure profiles embedded in environmental risk scenarios. TKTD simulations can also be linked with other types of models (*e*.*g*., Individual Based Model) within a pipeline of computing inference as for example Bayesian inference or Machine Learning. To handle such challenges within the particular framework of TKTD models for survival, we present an innovative simulation tool written in the new programming language Julia, called `TKTDsimulations.jl`. Given that TKTD models for survival usually require high performance computing due to the numerical integration of differential equations, our tool strongly benefits from Julia’s facilities, in particular a code that is fast to compile and easy to maintain. In addition, to ease the link with the already developed R-package `morse` dedicated to the statistical handling of ecotoxicity data, we also developed a new R-package, called `tktdjl2r`, interfacing `morse` with our new simulation tool `TKTDsimulations.jl` that considerably faster predictions with the corresponding ready-to-use `morse` functions.

## 1 Introduction

Toxicokinetics-Toxicodynamics (TKTD) models are increasingly used for the inference of toxicity indices of interest in Environmental Risk Assessment (ERA) because they can offer a clear description of numerous mechanisms, from the kinetics of compounds inside organisms (namely, the toxicokinetic, TK part) to their related damages and effect dynamics at the individual level (namely, the toxicodynamic, TD part) (Jager & Ashauer 2018, EFSA 2018). TKTD models offer the advantage of accounting for temporal aspects of exposure and toxicity considering data points all along the experiments, so that no data are lost. And their conceptual background allows their use under untested situations with time-variable exposure profiles either measured in the field or simulated within environmental risk scenarios (Baudrot & Charles 2019). Given all these advantages, TKTD models have been promoted to conduct risk assessments that are recognized to be more integrative, coherent and effective (OECD 2006, on Plant Protection Products & their Residues PPR, EFSA 2018). In particular, the General Unified Threshold model of Survival (GUTS) which have been proposed to unify most of the TKTD models for survival (Jager et al. 2011, Jager & Ashauer 2018) is today recognized and supported as ready-to-use for ERA (EFSA 2018).

We previously worked on Bayesian inference of GUTS models (Baudrot, Preux, Ducrot, Pave & Charles 2018) that we implemented within the R-package `morse`. Bayesian inference is tailored for decision making as it provides assessors with a range of values, namely the joint posterior distribution, that quantifies the uncertainty on parameters simultaneously, what is particularly valuable in risk assessment (Baudrot & Charles 2019). Indeed, the joint posterior distribution of parameters can be direclty used to predict survival curves under tested and untested exposure profiles, to calculate LC(x, t) and MF(x, t), and to compute goodness-of-fit measures (see hereinafter) (Baudrot & Charles 2019).

Based on this previous work on Bayesian inference, we present in this paper an innovative generic computing framework to make simple the use of GUTS, and its different versions, under complex exposure scenarios usually requiring high-performance computing (*e*.*g*., CPUs parallelization, GPUs). This kind of challenge is in fact not only specific to GUTS models, but also concerns models such as spatially-explicit agent-based models (Baudrot et al. 2021) or computational-intensive inference approaches (*e*.*g*., Bayesian (Baudrot, Preux, Ducrot, Pave & Charles 2018) and/or Machine-Learning).

Decision making in ERA requires the simulation of a great variety of environmental risk scenarios to be as realistic as possible, and as close as possible to the specificity of each landscape context. In this perspective, we implemented our new computing framework based on the recent language Julia, which “looks like Python, feels Like lisp, runs like Fortran” (Bezanson et al. 2017). Julia is designed to be well adapted for scientists requiring very fast simulation tools to link big data with complex models, as those that are more and more used and/or promoted within ERA. The Julia language strongly benefits of the `DifferentialEquations.jl`package for solving differential equations (Rackauckas & Nie 2017) which is the main dependence of our own `TKTDsimulation.jl` package. Hereafter, we thus present our GUTS models implementation in Julia. Then we provide an illustrative application based on standard toxicity test data.

## 2 Implementation of GUTS models

The GUTS modelling approach is a theoretical framework dealing with stressor effects on survival over time, based on hypotheses related to the stressor quantification, some possible compensatory processes (damage and/or recovery) and the nature of the death process (considered as a stochastic process for all organisms, or individually for each organism).

### 2.1 Toxicokinetics

In this paper, we use the simplest toxicokinetic (TK) writing of GUTS versions, with only one-compartment. The exposure profile *C*_*ext*_(*t*) can be either constant (*i*.*e*., independent on time *t*), or variable in the environment over time, leading to what we are classically calling an exposure profile. The internal concentration *C*_*int*_(*t*) within organisms consequently changes with time as the difference between uptake and elimination fluxes whose rate are denoted *k*_*u*_ and *k*_*e*_, respectively. A reduced form of the TK part of the model can be provided by introducing the ratio of both rates as a new parameter *k*_*u/e*_ = *k*_*u*_*/k*_*e*_ (also know as the bio-concentration factor, or BCF):

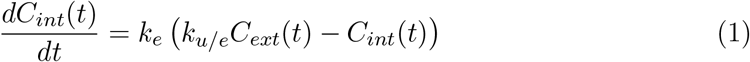

The external concentration or the exposure profile *C*_*ext*_(*t*) are experimentally measured at *n* time points *T* = *{T*_1_, *…, T*_*n*_*}* corresponding to *n* observations *C*_*ext*_(*T*) = *{C*_1_, *…, C*_*n*_*}*. Assuming that the concentration between two time points *T*_*i*_ and *T*_*j*_ is linear and zero outside leads to the following writing:

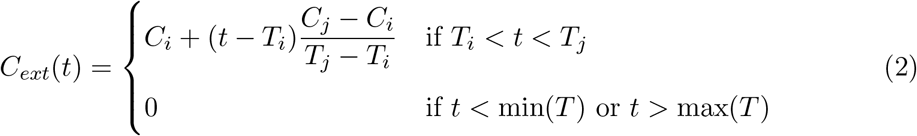

In Julia, equation (2) is implemented as follows:

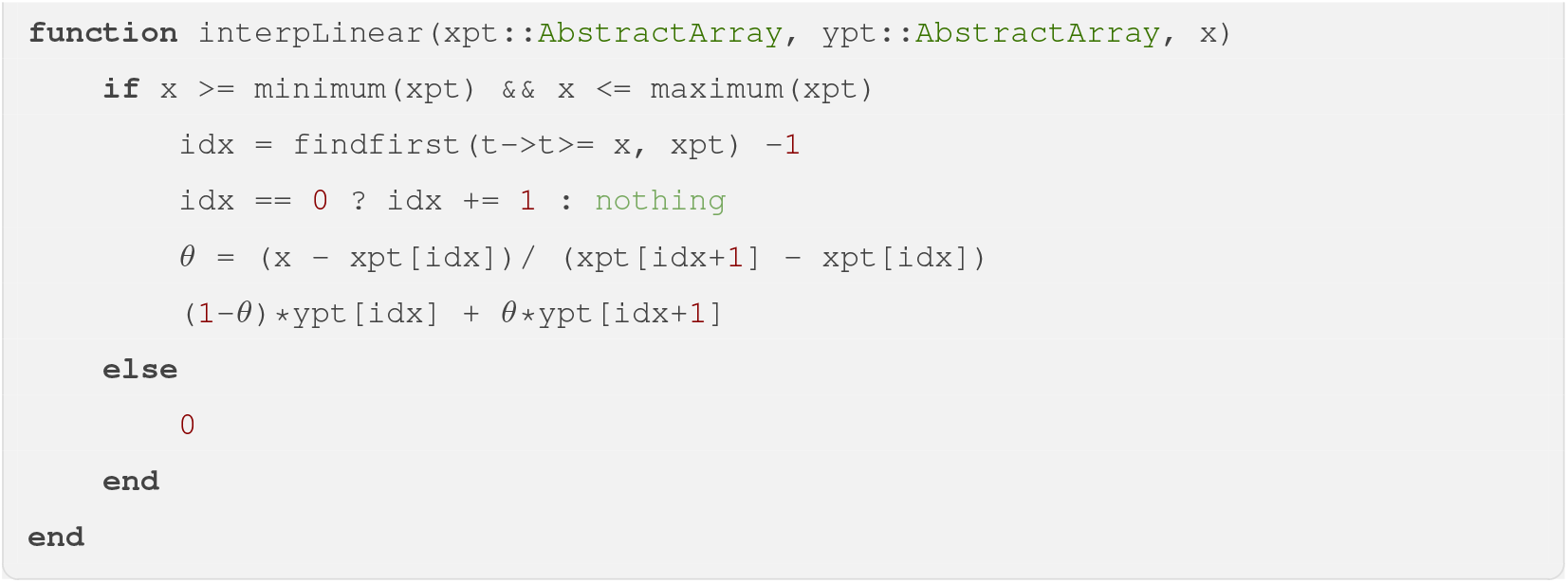

The internal concentration *C*_*int*_(*t*) induces damages within organisms, denoted with variable *D*(*t*). It is possible to describe the dynamics of *D*(*t*) with the one-compartmental approach of Jager & Ashauer (2018):

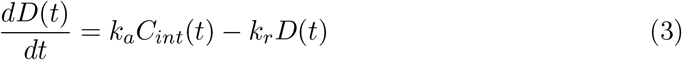

with *k*_*a*_ and *k*_*r*_ the rate constants of damage accrual and damage repair, respectively. By setting *k*_*d*_ as the ratio 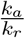 we get a normalized writing of equation (1), with the new parameter *k*_*d*_ that can be interpreted as the *dominant rate constant* (that is rate constant corresponding to the fastest event between damage accrual and damage repair):

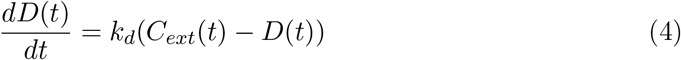

In Julia, equation (3) can be implemented in order to be solved by package `DifferentialEquation.jl`, leading to the following code:

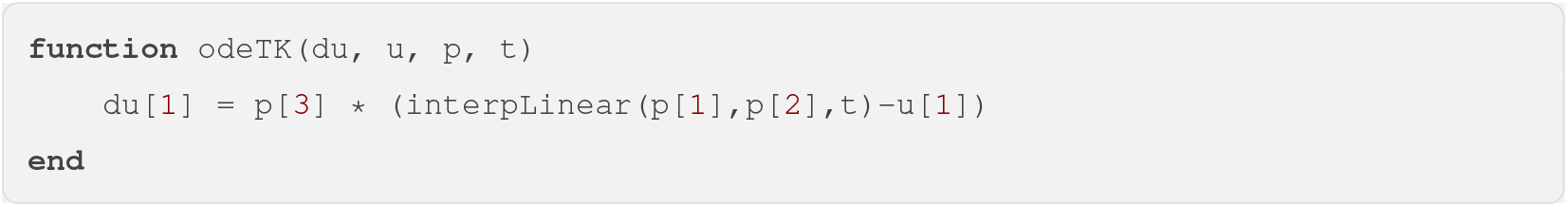

The final implementation of the whole TK model embeds the solver function as given in the code below:

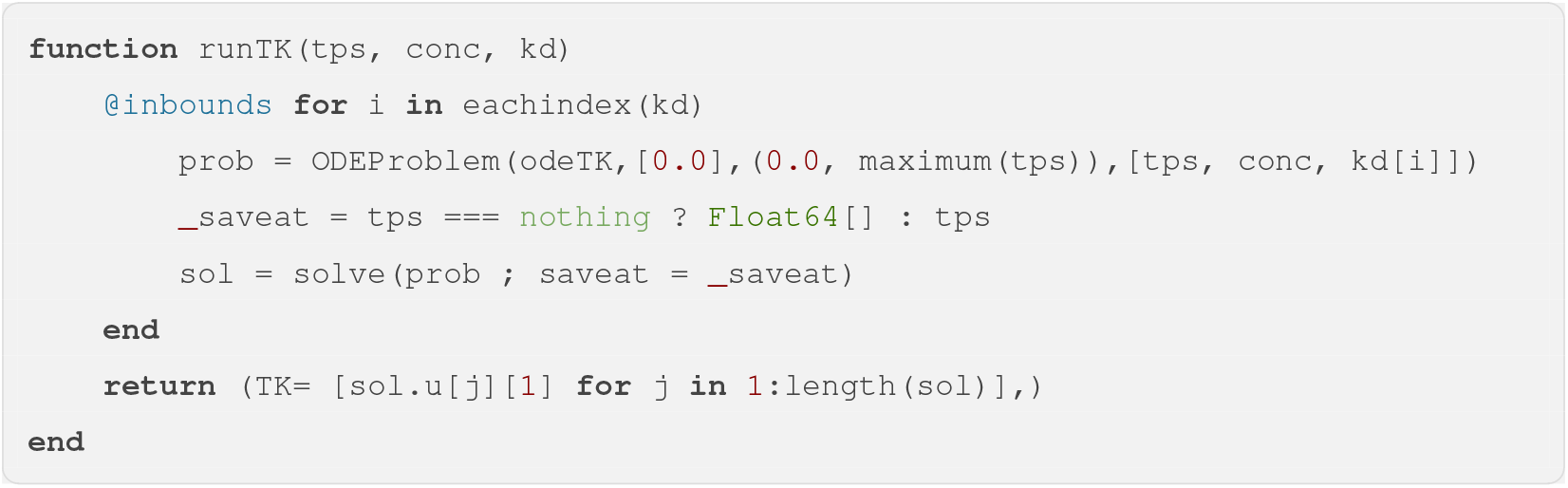

Let’s consider a classical FOCUS profile as given in Figure 1, that can be obtained as originally recommended by Boesten et al. (2000). See the corresponding Julia code below.

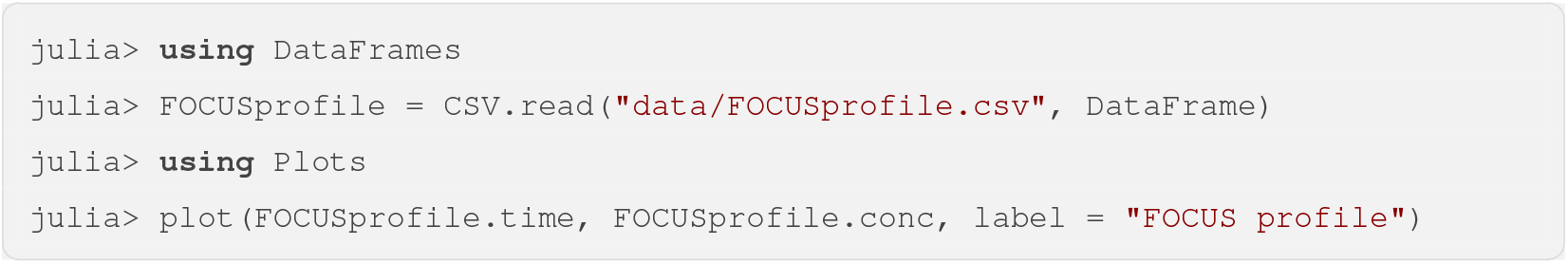

**Figure 1:**
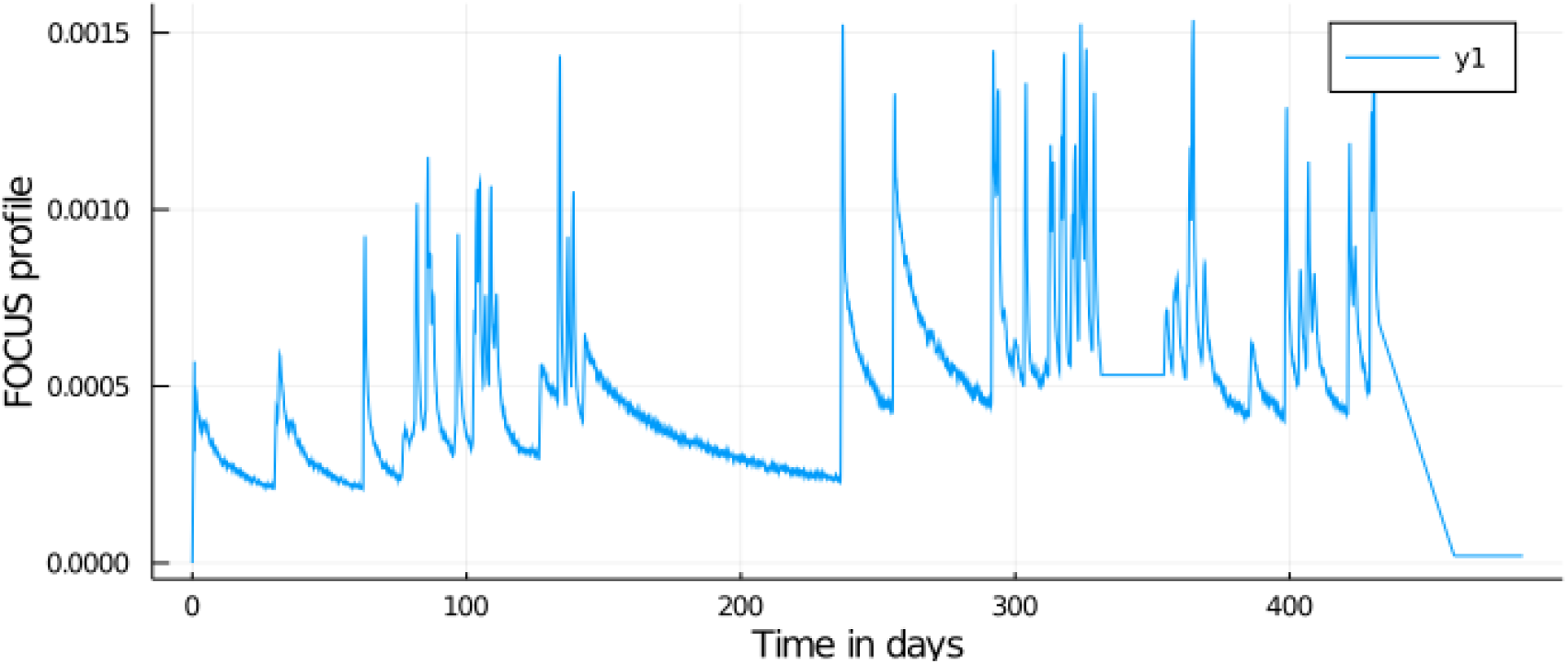
A classical FOCUS profile of exposure.

Running the TK part of the model in pure Julia can be performed according to the following code lines to get the predicted internal concentration over time:

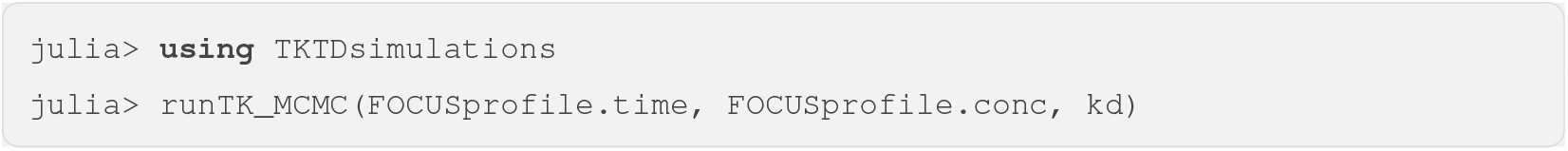

The previous command lines return as many TK time series as the dimension of *kd* parameter posterior distribution from which the predicted scaled internal concentration can be plotted with its uncertainty band (Figure 3).

The first run in Julia is longer than all the following ones because of the Just In Time Compilation process (Bezanson et al. 2017). However, from the second run, the speed with only one parameter becomes less than 0.5 seconds (see Figure 2). The same simulation can be performed with the R software by loading our new R-package `tktdjl2r` we specifically developed to make the interface with the Julia package. The command line `tktdjl2r::tktdjl2r_setup()` is required to make the link with `TKTDsimulations.jl` Julia package.

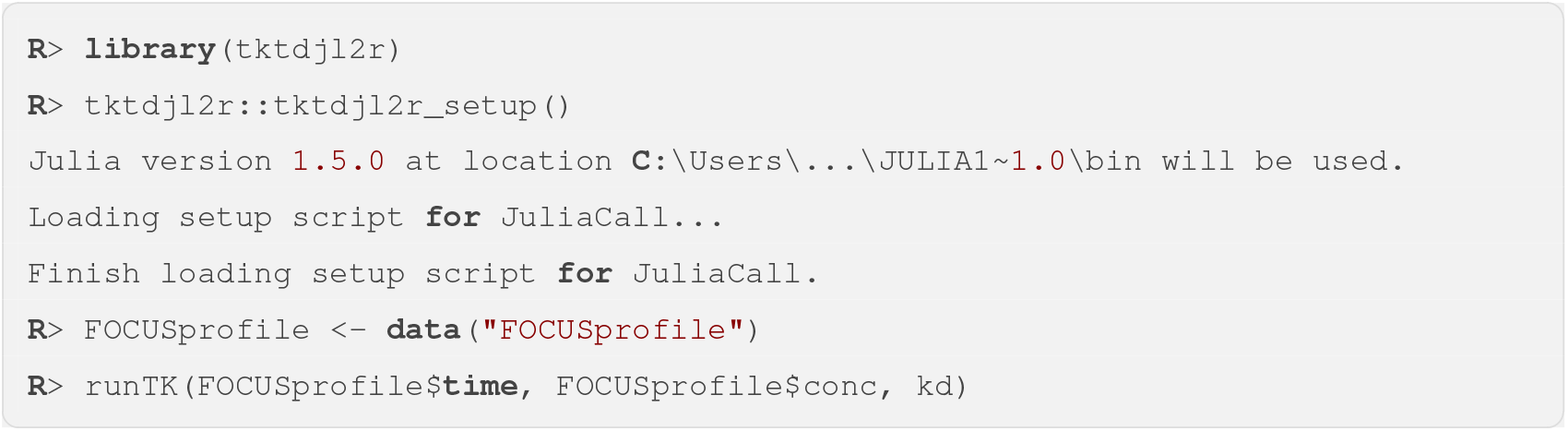

**Figure 2:**
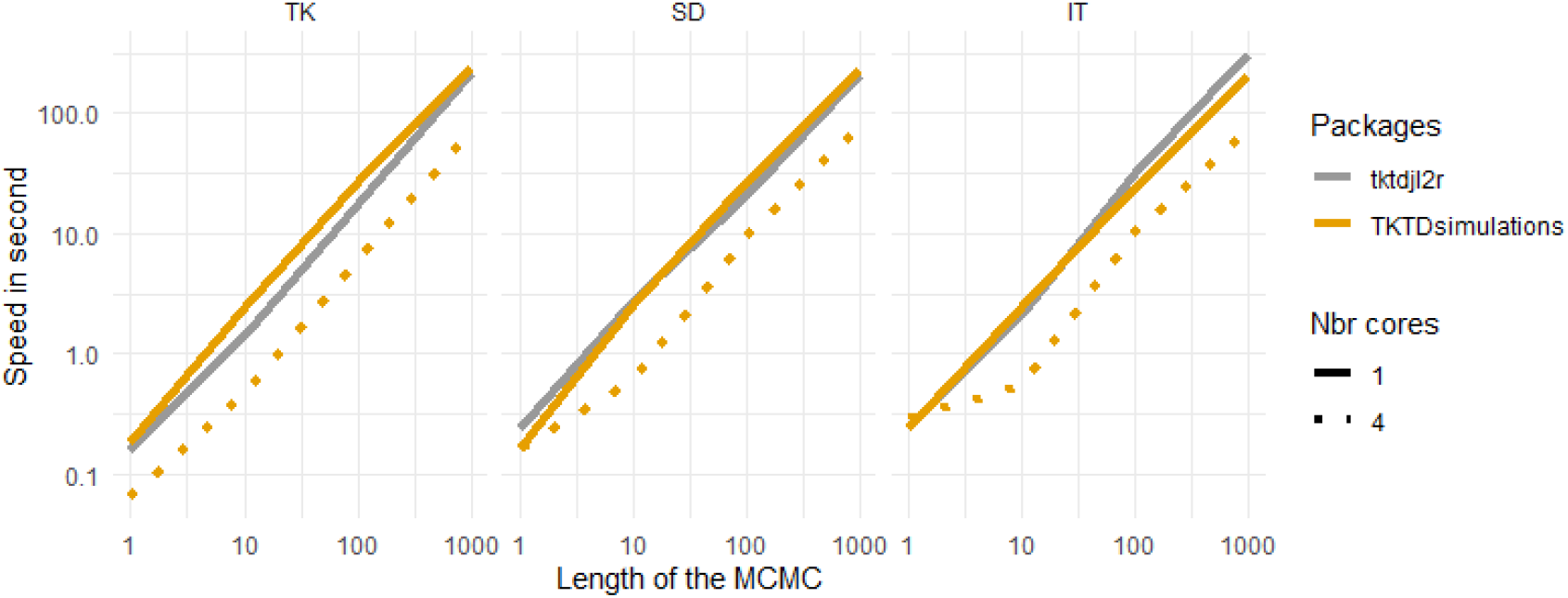
Speed of computing models TK, SD and IT depending on the length of MCMC chain using FOCUS profiles given by Figure 1.

**Figure 3:**
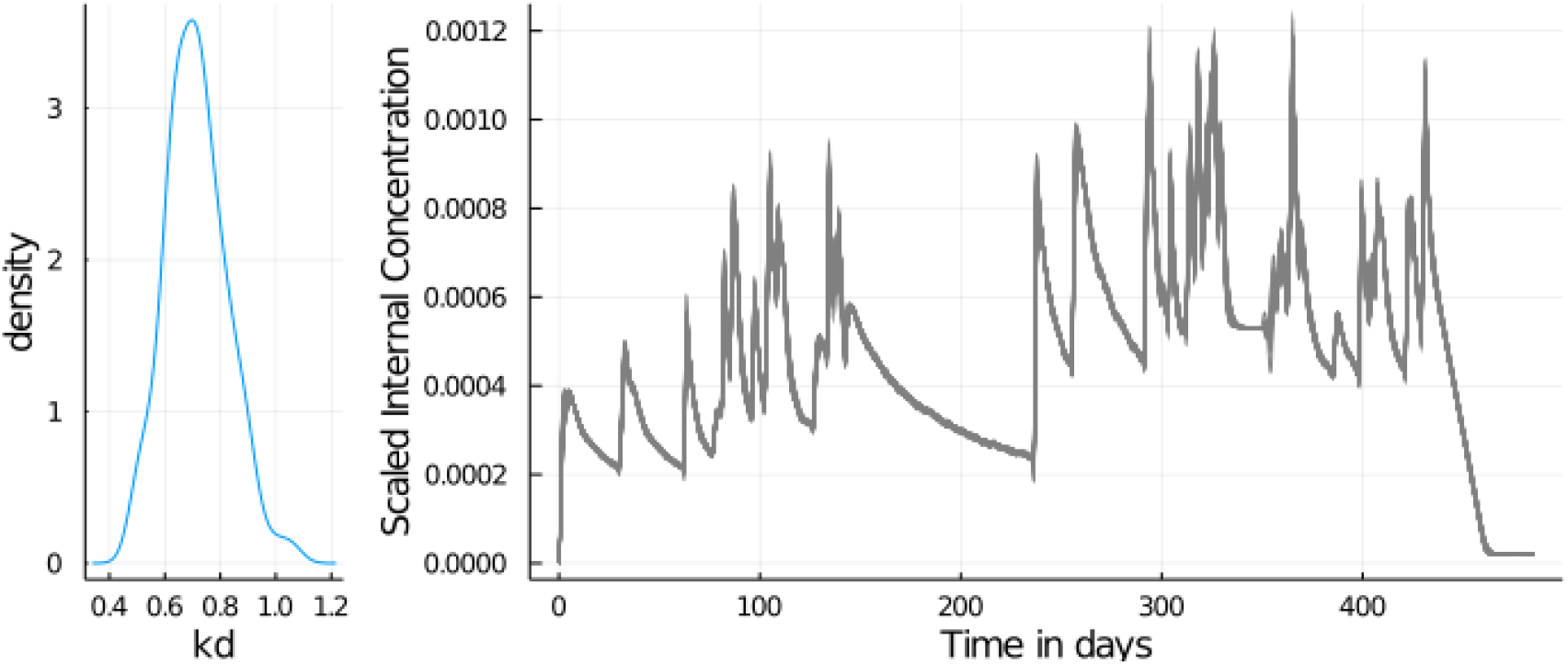
(left) distribution of the parameter *kd*. (right) output of the command line `runTK_MCMC()`.

Figure 2 shows that speed increases linearly with the number of time series (i.e., number of parameter estimates in the MCMC simulations), and that the use of Julia implementation from R is equivalent in term of speed. Of course, with a short MCMC sampling, calling the R-function is longer, since it requires almost 1 second to call Julia functions. However, this time is not going to increase if the data set is bigger since it is read only once. The simplicity of using multi-threading in Julia by increasing the number of cores to solve the ODEs thus increasing the speed of the calculations.

### 2.2 Toxicodynamics for survival

In the present paper, we consider the two most used derivations of GUTS models in their reduced form, namely both the stochastic death (GUTS-RED-SD) and the individual tolerance (GUTS-RED-IT) models (Jager & Ashauer 2018).

#### 2.2.1 Stochastic Death: GUTS-RED-SD

The GUTS-RED-SD model assumes that all individuals are identically sensitive to the chemical substance by sharing a common internal threshold concentration and that mortality is a stochastic process once this threshold is exceeded. Equation (5) describes the hazard rate as a function of the damage variable *D*(*t*) obtained from equations described in the toxicokinetic section above. Parameter *z* is the threshold parameter below which there is no instantaneous effect of the damage on the instantaneous hazard rate *h*_*z*_(*t*). Parameter *k*_*k*_ is the killing rate constant, corresponding to the slope of the linear function relating *h*_*z*_(*t*) with the damage variable *D*(*t*).

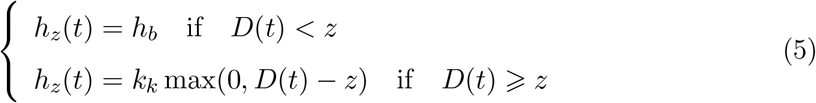

When damage is below threshold *z*, the instantaneous mortality rate simply equals to parameter *h*_*b*_ referring to the background mortality. The integration of the hazard rate function provides the survival probability until time *t*, denoted *S*(*t*) and given by:

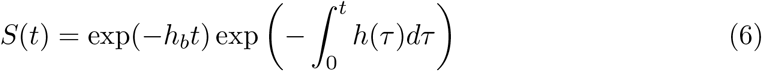

We can run the code in Julia with the following commands:

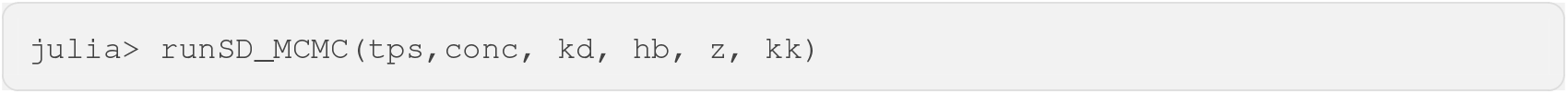

This command line return time series of survival probabilities from GUTS-RED-SD model as given by Figure 4. Using the R-package `tktdjl2r`, function *S*(*t*) for the GUTS-RED-SD model can be simulated based on equation (6) by simply using function `runSD` as illustrated below:

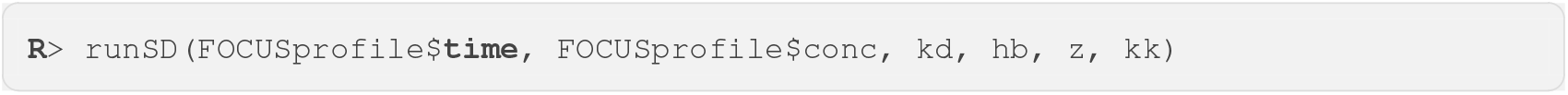

**Figure 4:**
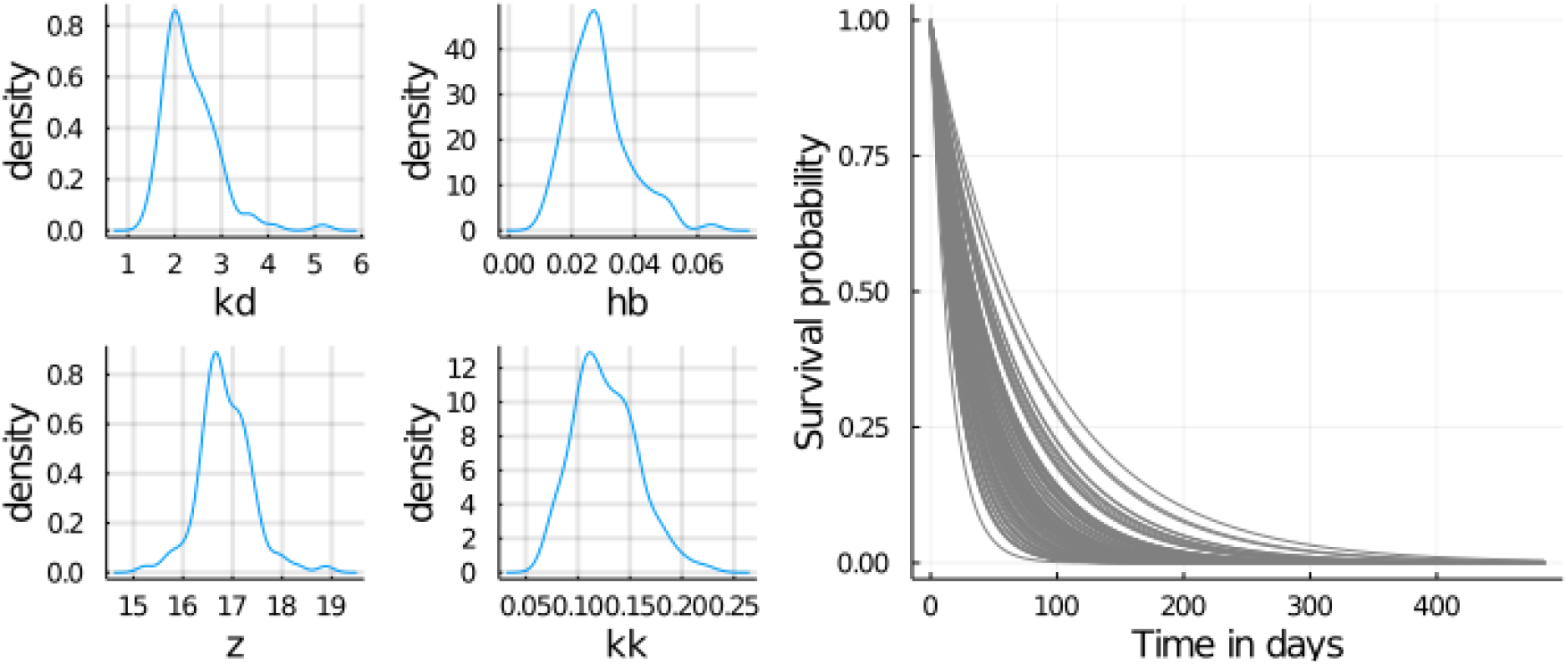
(left) distributions of the *k*_*d*_, *h*_*b*_, *z* and *k*_*k*_ parameters. (right) output of the command line `runSD_MCMC()` in Julia.

Both Julia and R simulation provide the same output of this command, the set of time series of TK and TD part of the GUTS-RED-SD model is illustrated by Figure 4.

#### 2.2.2 Individual Tolerance: GUTS-RED-IT

The GUTS-RED-IT model is based on the critical body residue (CBR) approach, which assumes that individuals differ in their tolerance threshold for a chemical compound according to a probability distribution. It also assumes that individuals die as soon as their internal concentration reaches their individual-specific threshold. Using the log-logistic probability distribution as the recommended default one for GUTS applications (Jager & Ashauer 2018), the median (*α*) and the shape (*β*) of the threshold distribution are the parameters of the toxicodynamic GUTS-RED-IT model in addition to the background mortality *h*_*b*_. Hence, the survival probability until time *t* can be written as follows:

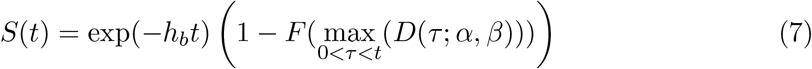

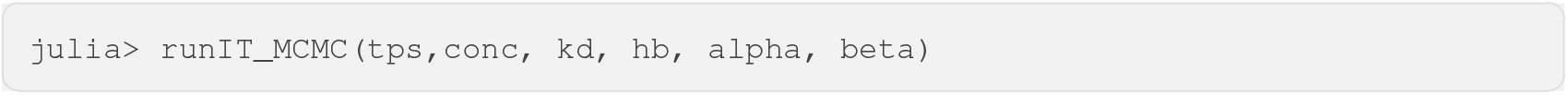

The command line above returns time series of survival probabilities from GUTS-RED-IT model as given by Figure 5.

**Figure 5:**
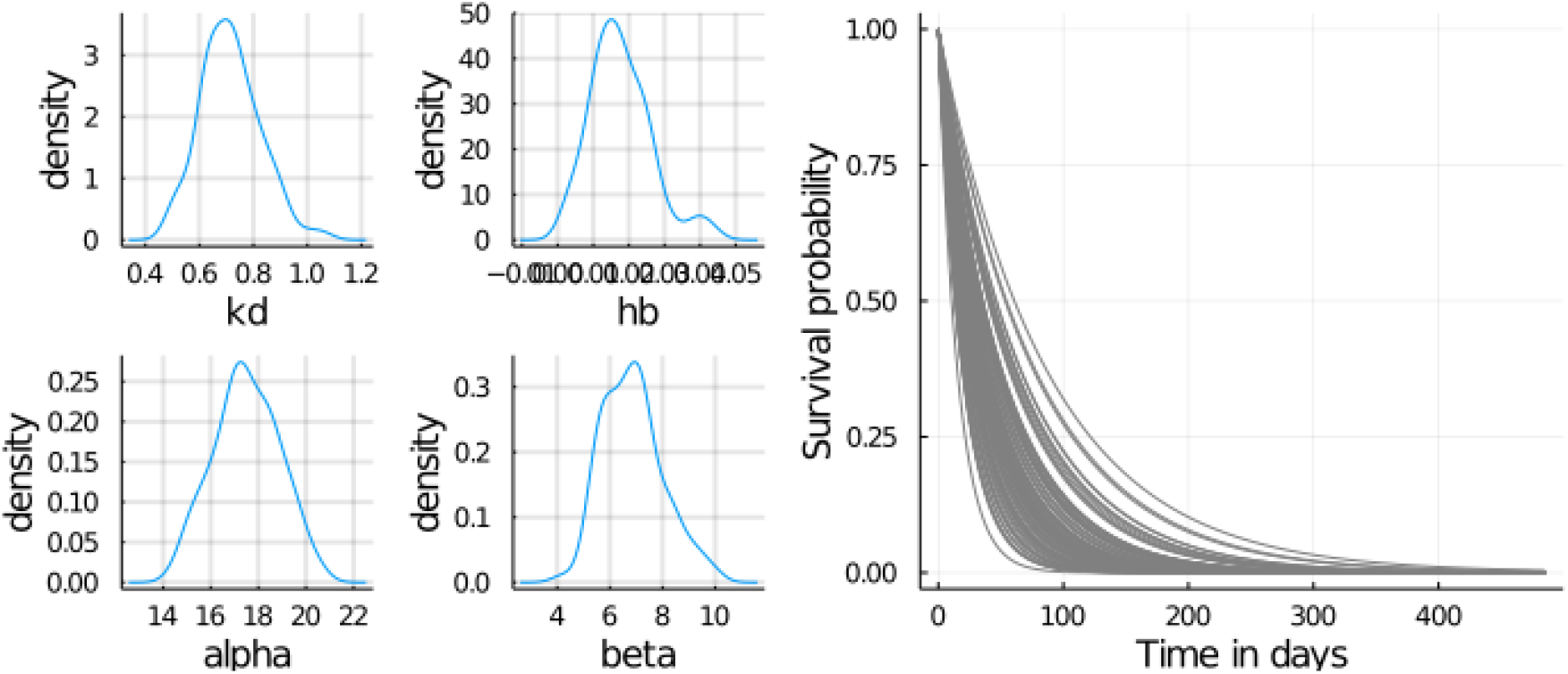
(left) distributions of the *k*_*d*_, *h*_*b*_, *α* and *β* parameters. (right) output of the command line `runIT_MCMC()` in Julia.

Using the R-package `tktdjl2r`, function *S*(*t*) for the GUTS-RED-IT model can be simulated based on equation (7) by simply using function `runIT` as illustrated below:

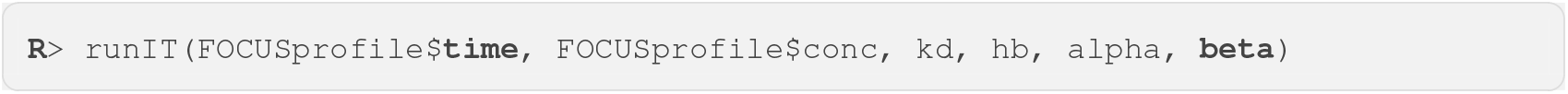

## 3 Application on standard data

In this last section, we illustrate our new simulation tool from the standard data set used in the EFSA scientific opinion (EFSA 2018). This data set concerns the lethal effects of propiconazole on the survival rate of the amphipod crustacean *Gammarus pulex* (Nyman et al. 2012). As recommended by EFSA (EFSA 2018), environmental risk assessment may be perform in a full workflow of three steps: (1-Calibration) inference of parameter estimates of both GUTS-RED models from experimental data collected under constant exposure conditions (as classically done in laboratory toxicity tests); (2-validation) the use of the previous parameter estimates to simulate the survival probability under a time-variable exposure profile specifically designed to also collect data. These data will be compared to the predictions based on validation criteria before going into the next step; (3-prediction) simulations with the most appropriate model regarding step (2) (namely the most conservative one) for environmentally realistic exposure scenarios of interest in a decision-making process. In this section, only steps (1) and (3) are illustrated.

First of all, the data set can be directly obtained from the R-package `morse` according to the following R code lines which provides Figure 6:

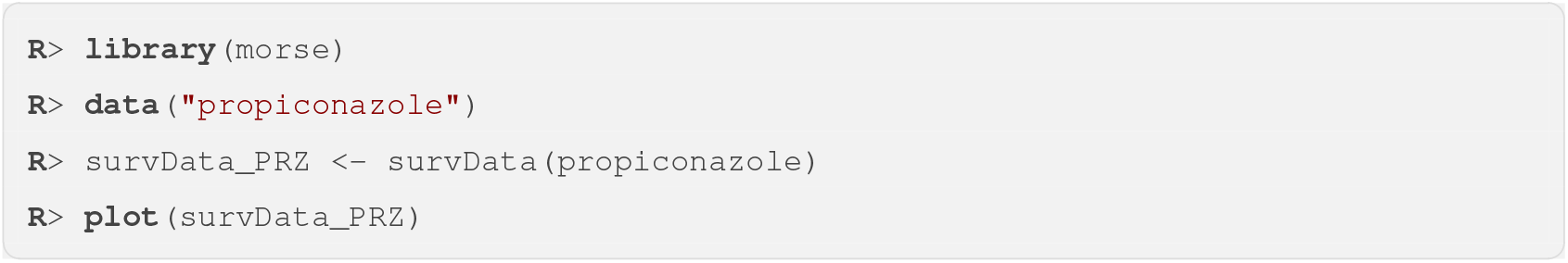

**Figure 6:**
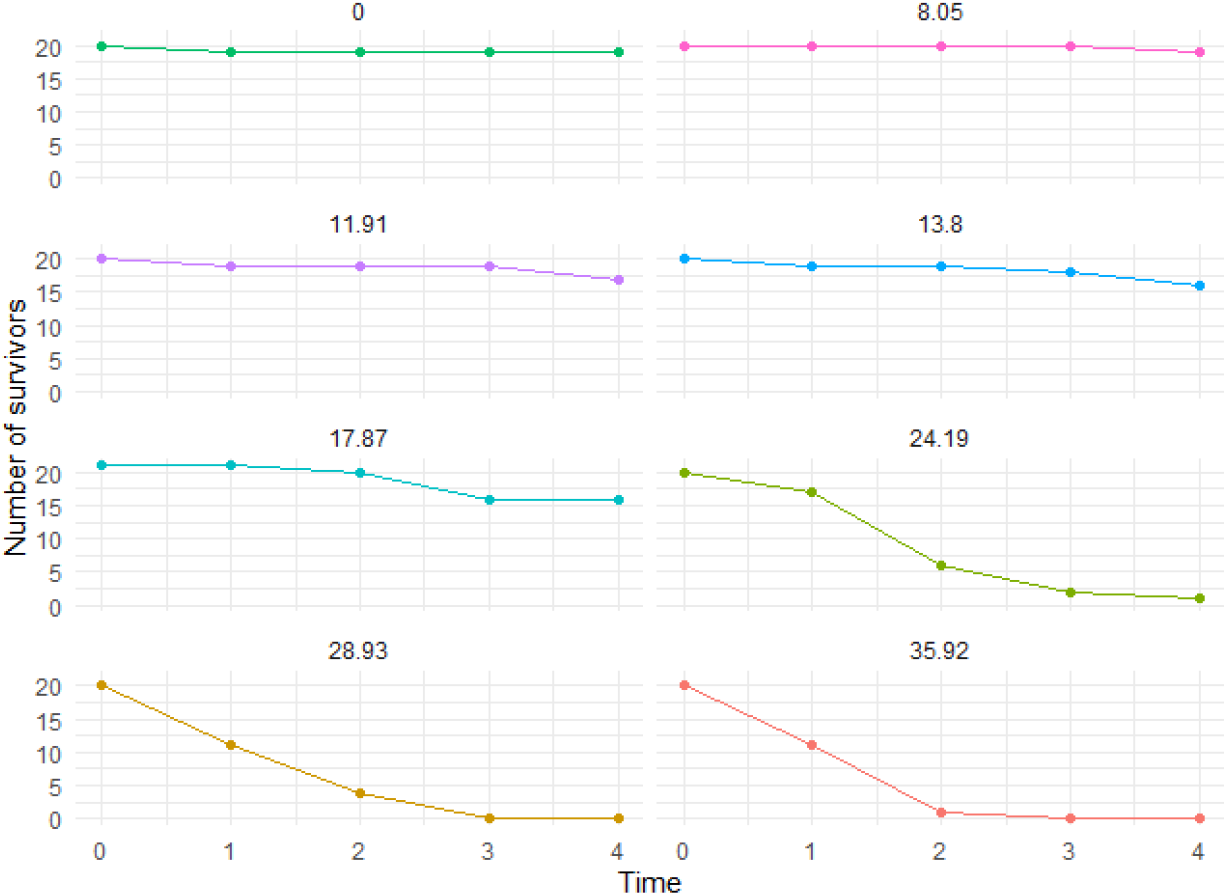
Results of experiments of survival of Gammarus pulex (Nyman et al. 2012).

Following the EFSA workflow, step (1) is then performed by fitting both GUTS-RED models to the loaded data set. As example, below are the R code lines for model GUTS-RED-SD fit with the Bayesian inference of the R-package `morse`:

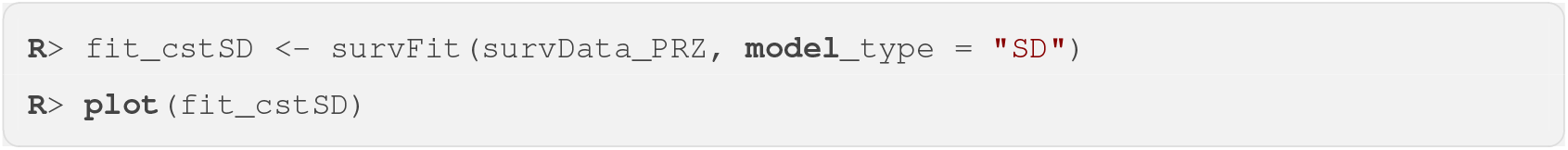

Please note that all Bayesian inference performed above in pure R language (basically by only using `morse`) can also be preformed directly on-line with the web platform MOSAIC, freely available here https://mosaic.univ-lyon1.fr/ (Baudrot, Veber, Gence & Charles 2018, Charles et al. 2018), from menu “surv”, then sub-menu “GUTS-fit” (EFSA workflow step (1)) or sub-menu “GUTS-predict” (EFSA workflow step (2) and (3)).

Outputs of the fitting object `fit_cstSD` are provided in Figure 7 where, for each of the tested concentrations, the median curve of the fit corresponds to the orange line, the 95% credible band to the grey area and the experimental data to the black dots associated with the observed 95% binomial confidence interval.

**Figure 7:**
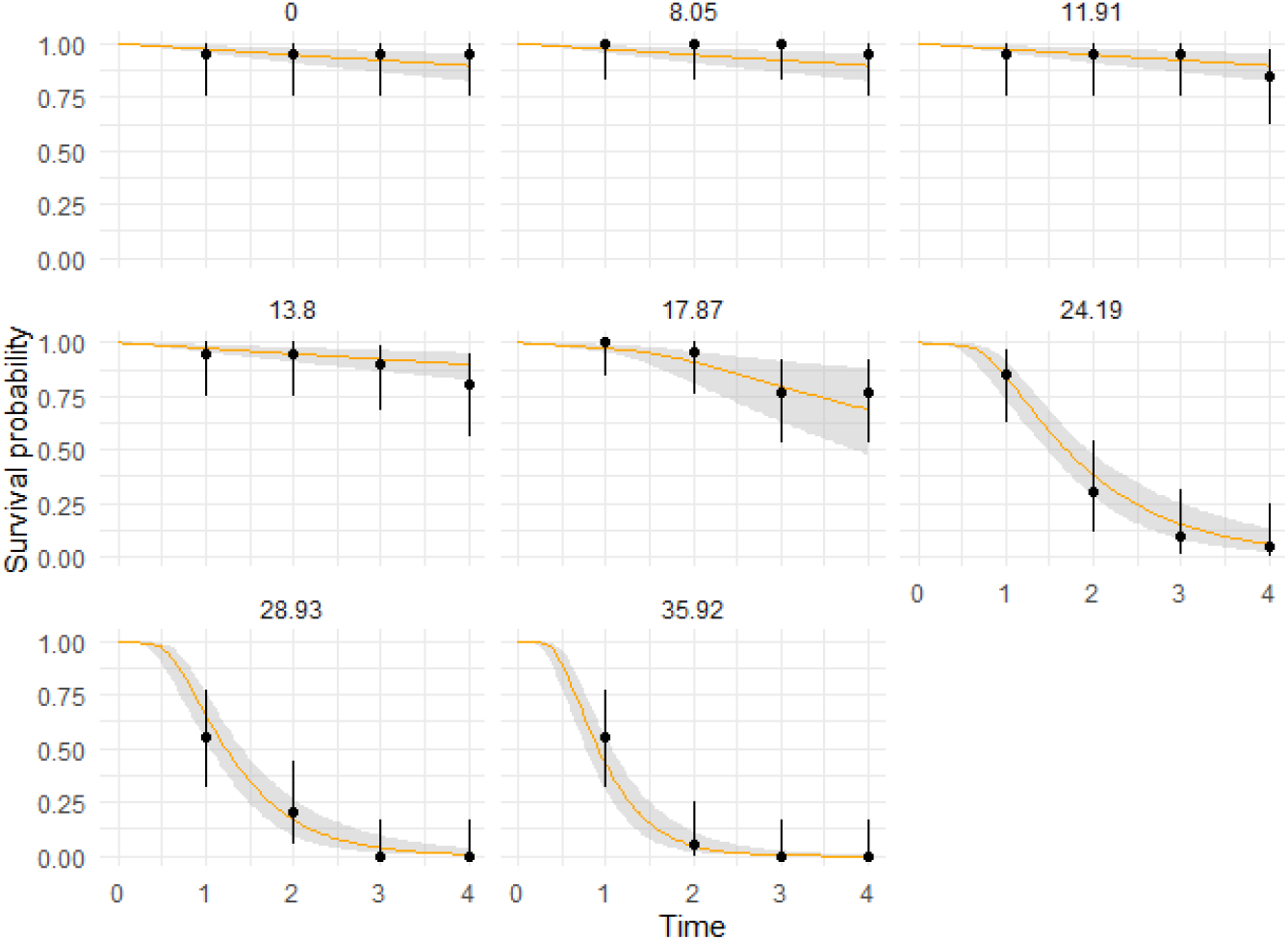
Fit results of data set provide in Figure 6 with the Bayesian inference package `morse`. Orange lines are estimated median and grey areas are 95% credible intervals.

The EFSA workflow step (3) can then be performed with the R-package `tktdjl2r` using command line given in Table 1, by simulating survival probability under either the GUTS-RED-SD or the GUTS-RED-IT model with parameter estimates as provided by the fitting objects obtained with the R-package `morse` (see step (1)).

**Table 1:** Functions exported by the R-package `tktdjl2r`.

| Function name | Description |
| --- | --- |
| `tktdjl2r_setup()` | loading `TKTDsimulations.jl` with R environment |
| `runTK` | solve toxicokinetics model |
| `runIT` | solve GUTS-RED-IT model |
| `runSD` | solve GUTS-RED-SD model |
| `runMFx` | solve GUTS-RED-IT or GUTS-RED-SD models to compute the multiplication factors. |

As example, fitting object `fit_cstSD` for the GUTS-RED-SD model returns 11238 samples of the four-parameters set, those samples being an approximation of the joint posterior probabilities for the four parameters of the model. Note that this number is not supposed to be exactly the same when performing again the simulations, as it relies on the stochasticity of the Monte Carlo Markov Chain algorithm underlying the Bayesian inference process as used in the R-package `morse`.

The FOCUS profile we used as example exposure profile to perform the simulations contains a total of 11641 time points. Below are the R code lines leading to the predictions of the survival probability over the entire profile as shown in Figure 8.

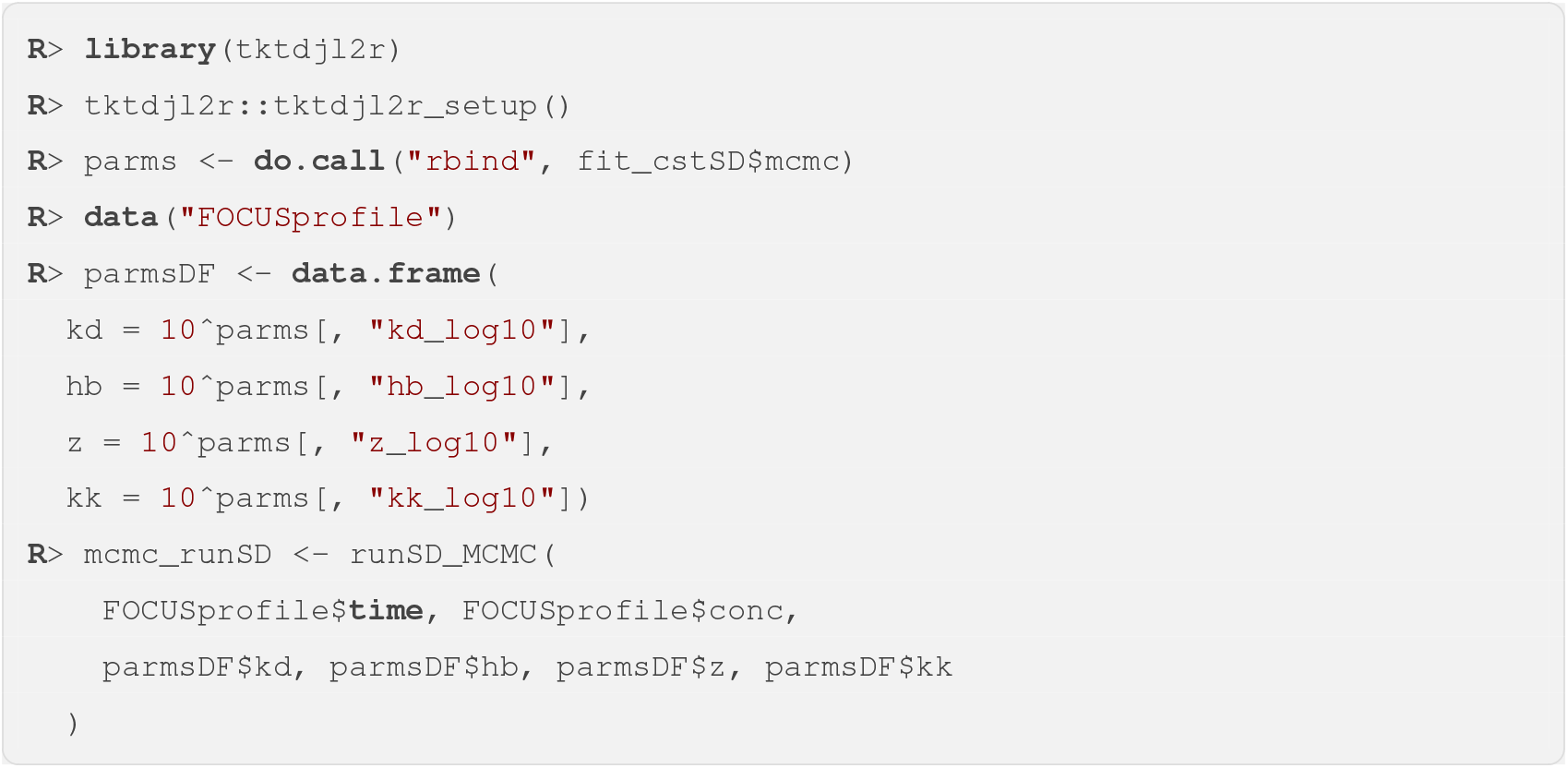

**Figure 8:**
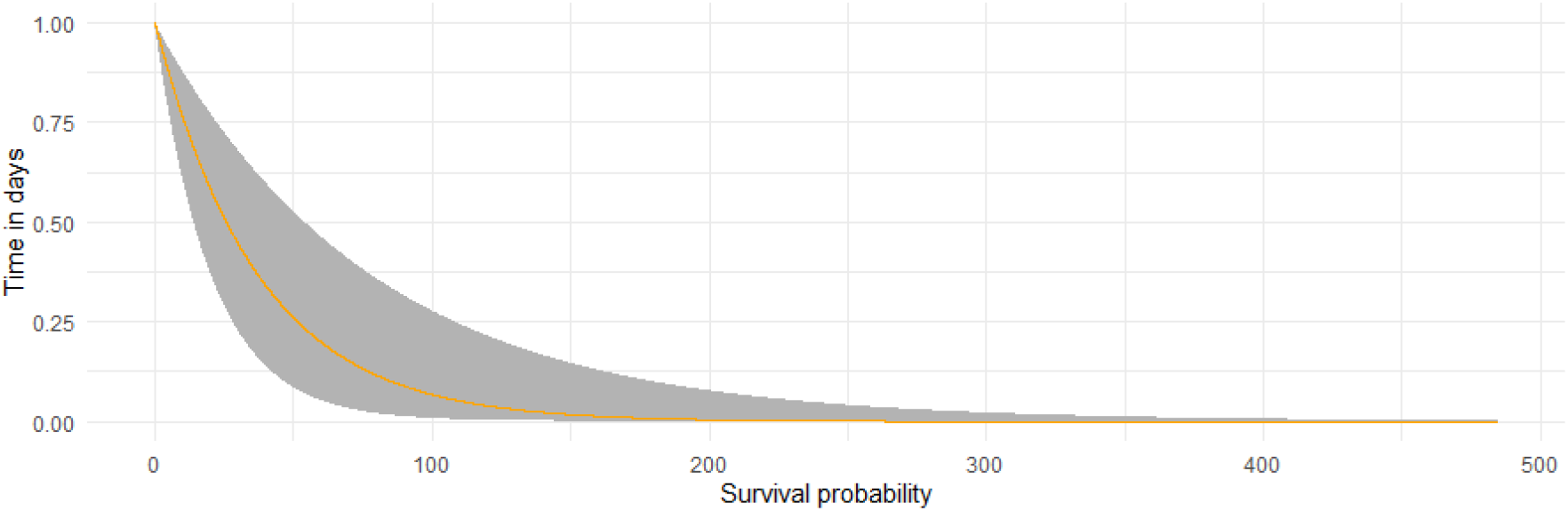
Prediction of the survival probability with the GUTS-RED-SD model using fitting object `fit_cstSD` plotted in Figure 7 over the FOCUS profile of Figure 1.

This basic example illustrates the straightforward link between Julia (namely, package `TKTDsimulations.jl`) and the R software (namely, packages `tktdjl2r` and `morse`) to serve as reference results for further users before performing they own analyses. Prediction results can easily be plotted by using classical R packages as given below for example (Figure 8).

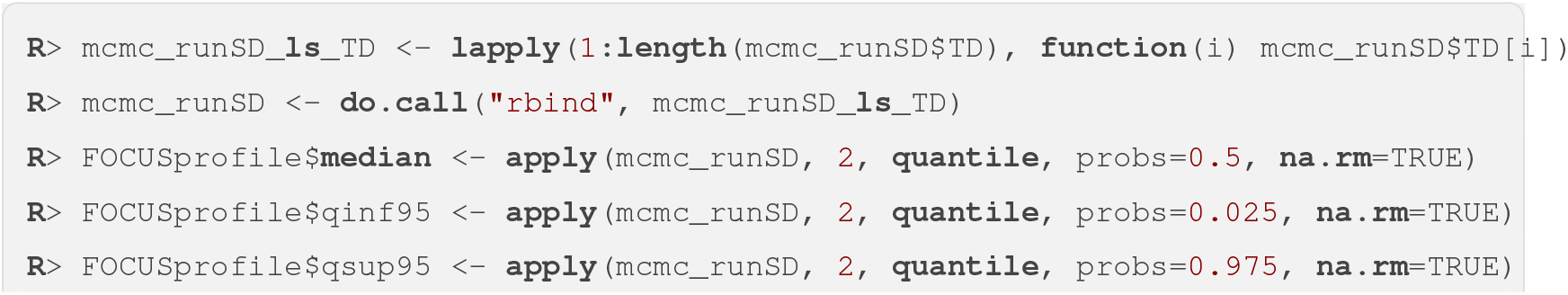

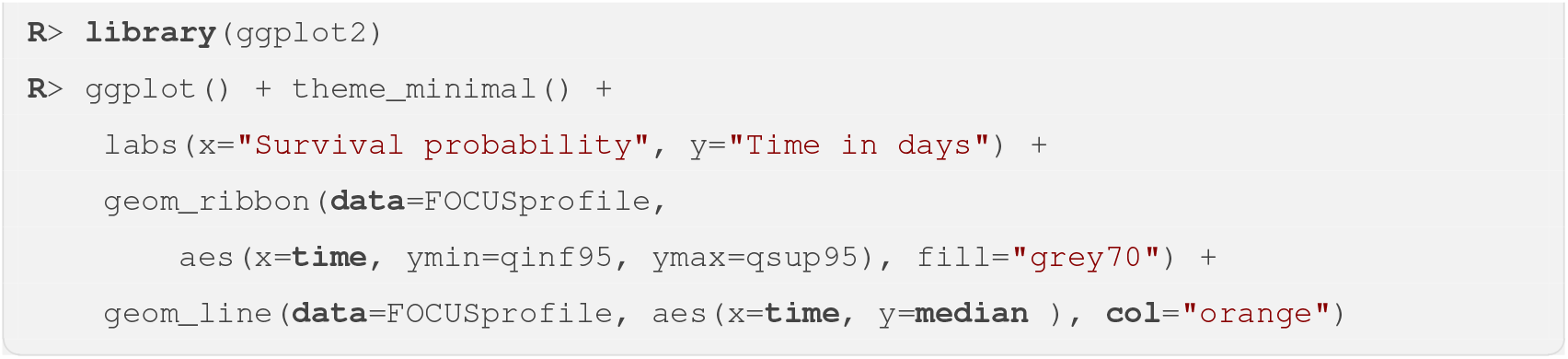

In environmental risk assessment, as firstly introduced by EFSA (EFSA 2018), the decision-making process also relies on the calculation of the maximum multiplication factor that could be applied to a given time-variable exposure profile according to an accepted *x*% additional reduction in the final survival probability, namely the calculation of the famous *x*% Lethal Profile (or *LP*_*x*_). This calculation can easily be performed with `morse` as given in the next section.

### 3.1 Computing Multiplication Factor

When applied to environmental exposure profiles, the Multiplication Factor gives the *x*% effect reduction at a specific time *t* (*MF* (*x, t*), or denoted *LP* (*x, t*) by EFSA) (Baudrot & Charles 2019). Considering a very simple exposure profile of only 7 time points (Figure 9) and using 11238 samples of parameters coming from the Bayesian inference using data of Figure 6 with GUTS-RED-IT model, the following commands line provides the specified *MF* (*x, t*) in 40.8 seconds.

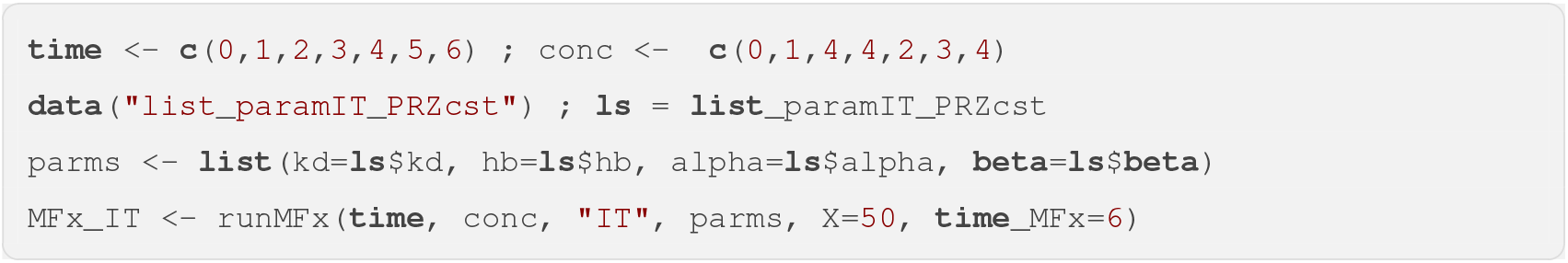

**Figure 9:**
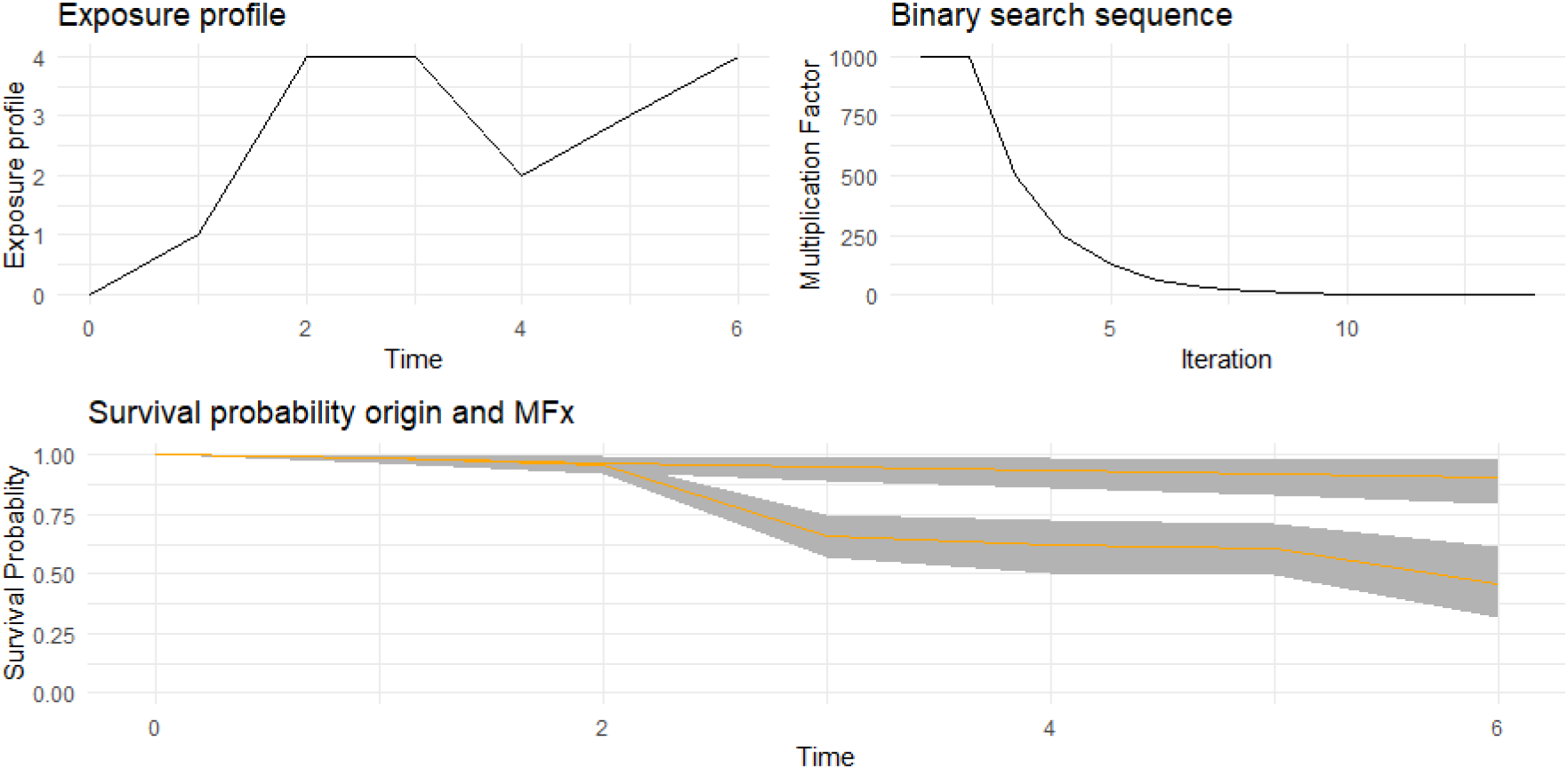
Multiplication Factor calculation leading to a reduction by 50% of the survival at target time *t* = 6 with GUTS-RED-IT model. (left-upper panel) exposure profile used, (right-upper panel) binary search sequences of *MFx* and (lower panel) survival time series from the original exposure profile and after application of the calculated Multiplication Factor *MFx*(50, 6).

The `MFx_IT` object returns the multiplication factor (here *MF* (*x* = 50, *t* = 6) = 5.61), the binary search sequence of the *MFx* as well as the TK and TD time series for both simulations, from the original exposure profile and after applying the calculated *MF*_*x*_ (Figure 9).

## 4 Discussion

GUTS models belong to the general TKTD mechanistic effect modelling approach that explicitly describes the toxicokinetic (TK) process, translating the external concentration of the chemical substance to which organisms are exposed in the medium, into an internal concentration or a latent variable accounting for damage. This TK part of the models is of real interest since this is usually not the external concentration that directly produces toxic effects, but rather complex metabolism processes with their own time dependencies (Jager & Ashauer 2018). The link between the internal chemical concentration and potential effects on life history traits is therefore fully time-dependent requiring a toxicodynamic (TD) part in the model to be also time-dependent.

The main advantage of fitting TKTD mechanistic effect models over statistical descriptions of experimental data is the possibility to extrapolate beyond the conditions of the experiment themselves (Baudrot & Charles 2019). The use of ecological risk scenarios that integrates spatially explicit exposure models with ecological effect models can then provide high-tier options for a prospective regulatory risk assessment (Franco et al. 2017). The challenge of applied ecotoxicology for risk-management decisions is to integrate ecotoxicity, life history and spatio-temporal distributions of both exposure and populations (Price & Thorbek 2014). To make the link from lab-experiment to risk-management decisions, Price & Thorbek (2014) identified three keys: i) the definition of protection goals matching societal requirements, ii) the development and use of spatially explicit exposure tools, and iii) the development of mechanistic effects models.

In the present paper, the used TKTD models (namely, GUTS-RED models) are the simplest case with only one-compartment model for the TK part and only survival as life history trait taken into account within the TD part. Our approach shows how simple it is to implement such models in Julia with the benefit of high speed computation, what would offer new perspectives to develop much more complex TKTD models in a near future.

## Acknowledgements

The authors are thankful to Ibacon for providing a financial support in developing these tools. This work benefited from the French GDR “Aquatic Ecotoxicology” framework which aims at fostering stimulating scientific discussions and collaborations for more integrative approaches. This work is part of the ANR project APPROve (ANR-18-CE34-0013) for an integrated approach to propose proteomics for bio-monitoring: accumulation, fate and multi-markers (https://anr.fr/Projet-ANR-18-CE34-0013).

This work was made under the umbrella of the Graduate School H2O’Lyon (ANR-17-EURE-0018) and “Université de Lyon” (UdL), as part of the program “Investissements d’Avenir” run by “Agence Nationale de la Recherche” (ANR). The authors are grateful to Dimitri MIKEC who contributed during his internship as a master-grade student from INSA Lyon.

